# Inflammatory Cytokines Can Induce Synthesis Of Type-I Interferon

**DOI:** 10.1101/2024.01.08.574713

**Authors:** Nikhil Sharma, Xu Qi, Patricia Kessler, Ganes C. Sen

## Abstract

Type I interferon (IFN) is induced in virus infected cells, secreted and it inhibits viral replication in neighboring cells. IFN is also an important player in many non-viral diseases and in the development of normal immune cells. Although the signaling pathways for IFN induction by viral RNA or DNA have been extensively studied, its mode of induction in uninfected cells remains obscure. Here, we report that inflammatory cytokines, such as TNF-α and IL-1β, can induce IFN-β through activation of the cytoplasmic RIG-I signaling pathway. However, RIG-I is activated not by RNA, but by PACT, the protein activator of PKR. In cell lines or primary cells expressing RIG-I and PACT, activation of the MAPK, p38, by cytokine signaling, leads to phosphorylation of PACT, which binds to primed RIG-I and activates its signaling pathway. Thus, a new mode of type I IFN induction by ubiquitous inflammatory cytokines has been revealed.

**Key points:**

- Cytochalasin D followed by TNF-α / IL-1β treatment activates IFN-β expression.
- IFN-β expression happens due to activation of RIG-I signaling.
- Interaction between RIG-I and PACT activates IFN-β expression.

## INTRODUCTION

In mammals, a major component of host defense to virus infection is mediated by type I interferons (IFN) (1). The signaling pathways used by viruses to induce IFN have been extensively studied; they are triggered primarily by recognition of intracellular viral nucleic acids by endosomal or cytoplasmic pattern recognition receptors such as TLRs, cGAS/STING or RLRs (2). However, IFNs are involved in many physiological and pathological processes, above and beyond viral infection. For example, B cell maturation and T cell functions require the participation of IFNs (3–5) and type I IFNs play both positive and negative roles in a variety of cancers and autoimmune diseases (6, 7). It is unclear how IFN is induced in the absence of virus infection, although cellular nucleic acids may participate in the process (8, 9).

Viral double-stranded (ds) RNA triggers several antiviral responses in an infected cell. It can activate RIG-I signaling to induce IFN, which, in turn, induces antiviral genes (10). The dsRNA can also activate two antiviral enzymes 2-5 OAS and PKR (11, 12). The dsRNA binding to PKR causes its activation by auto-phosphorylation and active PKR can inhibit protein synthesis by phosphorylating the translation initiation factor eIF-2α (12). An alternative activator of PKR is PACT, another dsRNA-binding protein, which can activate PKR without any involvement of dsRNA (13). The dsRNA-binding domains and the PKR-activation domain of PACT are distinct and two serine residues in the latter domain, S246 and S287, need to be phosphorylated for its ability to activate PKR in cells (14). S246 is constitutively phosphorylated but S287 is phosphorylated in response to various extracellular stresses (14). PACT has diverse physiological roles including its critical functions in anterior pituitary development in mice (15). Kok *et al* reported that PACT can facilitate RIG-I signaling in virus-infected cells (16). PACT can bind to the C-terminal domain of RIG-I in infected cells and augment its ability to signal (16). However, they did not observe endogenous RIG-I activation by endogenous PACT in uninfected cells.

Here we report that inflammatory cytokines, such as TNF-α or IL-1β, can induce IFN-β by activating PACT, which binds to RIG-I and trigger its signaling in the absence of any viral RNA. However, RIG-I needs to be primed for PACT activation. Acharya *et al* reported that viral activation of RIG-I is a two-step process; first RIG-I is dephosphorylated by a phosphatase and then the primed RIG-I is activated by viral dsRNA (17). The current study shows that PACT, activated by the MAP kinase, p38, can substitute for dsRNA to activate signaling by primed RIG-I and thereby provides a means to IFN production by uninfected cells exposed to inflammatory cytokines.

## MATERIALS AND METHODS

### Reagents and antibodies

Anti-PACT (ab-31967) antibody was purchased from Abcam and Anti-RIG-I (AG 20B-0009-C100) from Adipogen life sciences. K-63 linkage specific polyubiquitin antibody (5621S) and anti-actin (3700) were purchased from Cell Signaling technology (CST). Anti-TRIM-25 antibody (sc-166926) from Santa Cruz biotechnology. Anti-FLAG antibody (SAB4200071) was purchased from Millipore Sigma. Mouse IgG (sc-2025) and rabbit IgG (sc-2027) control antibodies were purchased from Santa Cruz biotechnology. Anti-FLAG conjugated agarose beads (A4596, Millipore Sigma) and Dynabeads protein A (10001D, Thermo Fisher scientific) were used for immunoprecipitation. For reconstitution of PACT (MG52537-NF) cDNA clone expression plasmid was purchased from Sino Biologicals. The WT PACT expression vector was used as template for constructing p38 mutant PACT clone by using Q5 site-directed mutagenesis kit (New England Biolabs, E0554S). Lipofectamine 2000 (Thermo Fisher Scientific, 11668030) was used for all transfection experiments. The clones were transfected into AML-12 cells and antibiotic selection was used to generate stable cell lines. Cytochalasin D (C8273, Neta) along with recombinant Mouse TNF-α (# RMTNFAI, Thermo Fisher Scientific) and human TNF-α (16769S, Cell Signaling technologies) were used for treatment. Mouse IL-1β was ordered from R and D systems (401-ML-005/CF). Mouse Interferon-β ELISA kit was obtained from PBL assay biosciences (42410–1). MAPK activator TPA / PMA (P1585 Sigma) and p38 inhibitor Adezmapimod (SB203580) (catalog no. S1076, Selleckchem.com) were used.

### Cell culture

Mouse AML-12 cell lines were maintained in DMEM-F12K 1:1 media (obtained from Cleveland Clinic Cell Culture services) supplemented with 10% FBS, 100 units/ml of penicillin, 100 mg/ml of streptomycin, 1x dexamethasone 10008980, Cayman chemicals), 1x insulin-selenium-transferrin (41400045, Thermo Fisher scientific). HepG2 cells and other cell lines were maintained in DMEM supplemented with 10% FBS, 100 units/ml of penicillin, and 100 mg/ml of streptomycin. RIG-I and PACT minus cells were generated in the lab as described preciously (18).

### mRNA quantification

MiRNeasy mini kit (Qiagen) was used for RNA isolation according to manufacturer’s protocol. cDNA synthesis was done using reverse transcriptase kit (#4368813, Thermo fisher scientific). For real-time PCR, 384 well-format real time PCR in a Roche Light Cycler 480 II using SYBR Green PCR reagents (#4309155, Thermo Fisher scientific). The primers used for mouse and human IFN-β mRNAs have been described before (18).

### Immunoprecipitation and Western blotting

These procedures were done as described before (18).

### Virus infection

Vesicular Stomatitis Virus (VSV) infection was given to cells at MOI 5 for one hour in serum free media. After 1 hour, serum free media was replaced by normal media.

### Proximity ligation assay (PLA)

Cells on glass slides were fixed using 4% paraformaldehyde and permeabilized with 0.25% Triton X-100, followed by blocking with 5% goat serum at room temperature. Subsequently, the cells were incubated overnight with primary antibodies against PACT (Abcam) and RIG-I (AdipoGen) in 5% goat serum. After washing, oligonucleotides-conjugated secondary antibodies, PLA Probe DUOLINK Anti-Mouse MINUS (DUO92004, Sigma) and PLA DUOLINK Probe Anti-Rabbit PLUS (DUO92002, Sigma), were used for incubation at 37°C for 1h. Cells were then incubated in the presence of DNA ligase for 30 min at 37°C. Finally, PLA signals were amplified by incubating cells with DNA polymerase for 100 min at 37°C (DUOLINK in Situ detection reagent red, DUO92008, Sigma). Confocal images were acquired using a Leica laser scanning confocal microscope and analyzed via Fiji software as previously described (19).

### Cytochalasin D, TNF-**α**, IL-1**β**, TPA and p38 inhibitor treatments

Cells were pre-treated with cytochalasin D (0.5µg/ml) for one hour followed by TNF-α or IL-1β treatment (100ng/ml). The cells were then harvested 1 hour post TNF-α or IL-1β treatment.

For p38 inhibitor experiments, 1µM of inhibitor was added to cells (1hr) before cytochalasin D or cytochalasin D + TNF-α treatment.

For MAPK activation, TPA (12-O-Tetradecanoylphorbol-13-acetate) treatment was given to AML-12 cells. DMSO or p38 inhibitor pre-treatment was given prior TPA treatment to cells to check the efficacy of p38 MAPK inhibitor.

## ELISA

For ELISA, culture supernatants were collected at 10 hours post treatment. Interferon-β was measured by ELISA kit (BD biosciences) following manufacturers’ instructions.

### PACT mutant generation

PACT Y305, at the putative p38 MAPK binding site (F-site), 18 amino acid residues downstream of S287 was mutated to Ala using site directed mutagenesis kit following manufacturer’s protocol. The double phosphomimetic mutant S246D, S287D (DD mutant) of PACT was previously constructed in our lab (ref).

The mutant PACT constructs were transfected into AML-12 cells with PACT minus background and cells were selected using puromycin selection marker to generate stable cell lines.

Primers used for site directed mutagenesis-

Y305A forward: 5’ TGC TCT GCA GGC GTT AAA GAT AAT AGC AG 3’

Y305A reverse: 5’ TTG TGA GCA GCG TC 3’

### Statistical analysis

All statistical analyses were performed using GraphPad Prism 9.5.1 software. The mean ± SEM of all biological replicates was used to make graphs. Statistical significance was calculated by student’s T-test or by one way ANOVA followed by turkey’s post hoc test. (P value ≤ 0.05 was considered significant, *P < 0.05; **P < 0.01, ***P < 0.001).

## RESULTS AND DISCUSSION

We have recently reported that in the mouse liver cell line AML-12, RIG-I mediated viral induction of IFN-β requires the presence of PACT (18). Like PKR activation, domain 3 of PACT is required to activate RLR signaling and substitution of two Serine residues by Ala in this domain, S246 and S287, abolished PACT’s ability to support viral induction of IFN (18). In contrast, their substitution by Asp, a mimetic of phospho-serine, enhanced IFN induction by VSV infection (Fig 1A). Surprisingly, the S246D, S287D mutant (DD) of PACT could induce IFN-β without virus infection. This led us to explore the possibility that activation of endogenous PACT by extracellular stresses may cause IFN induction. However, treating cells with known PACT activator, such as tunicamycin or actinomycin D, failed to induce IFN synthesis. Acharya *et al* reported that RIG-I activation is a two-step process: binding of viral RNA follows its prior priming through dephosphorylation, which requires the release of a subunit of the relevant phosphatase from the cytoskeletal actin filaments to the cytoplasm (17). Virus infection, transfection reagents or known cytoskeleton disruptors, such as cytochalasin D (Cyt D), all can provide the priming signal. We speculated that active PACT may trigger RIG-I signaling, but only if it is primed. Indeed, priming RIG-I by pre-treating cells with cytochalasin D, enabled PACT activators to induce IFN. IFN-β mRNA was strongly induced by TNF-α in Cyt D-treated cells (Fig 1B). The same was true for IL-1β (Fig 1C). As expected, IFN-β protein was secreted by the cells treated with TNF-α, but only if they were primed. (Fig 1D). IFN induction by TNF-α required the presence of both RIG-I and PACT, because ablation of expression of either protein abolished IFN-β induction (Fig 1E). The phenomenon was not cell-type specific or species-specific; several mouse and human cell lines, that expressed RIG-I and PACT, were responsive to TNF-α to induce IFN-β (data not shown). Importantly, primary cells, such as mouse hepatocytes, responded similarly (Fig 1F).

**Figure 1.**
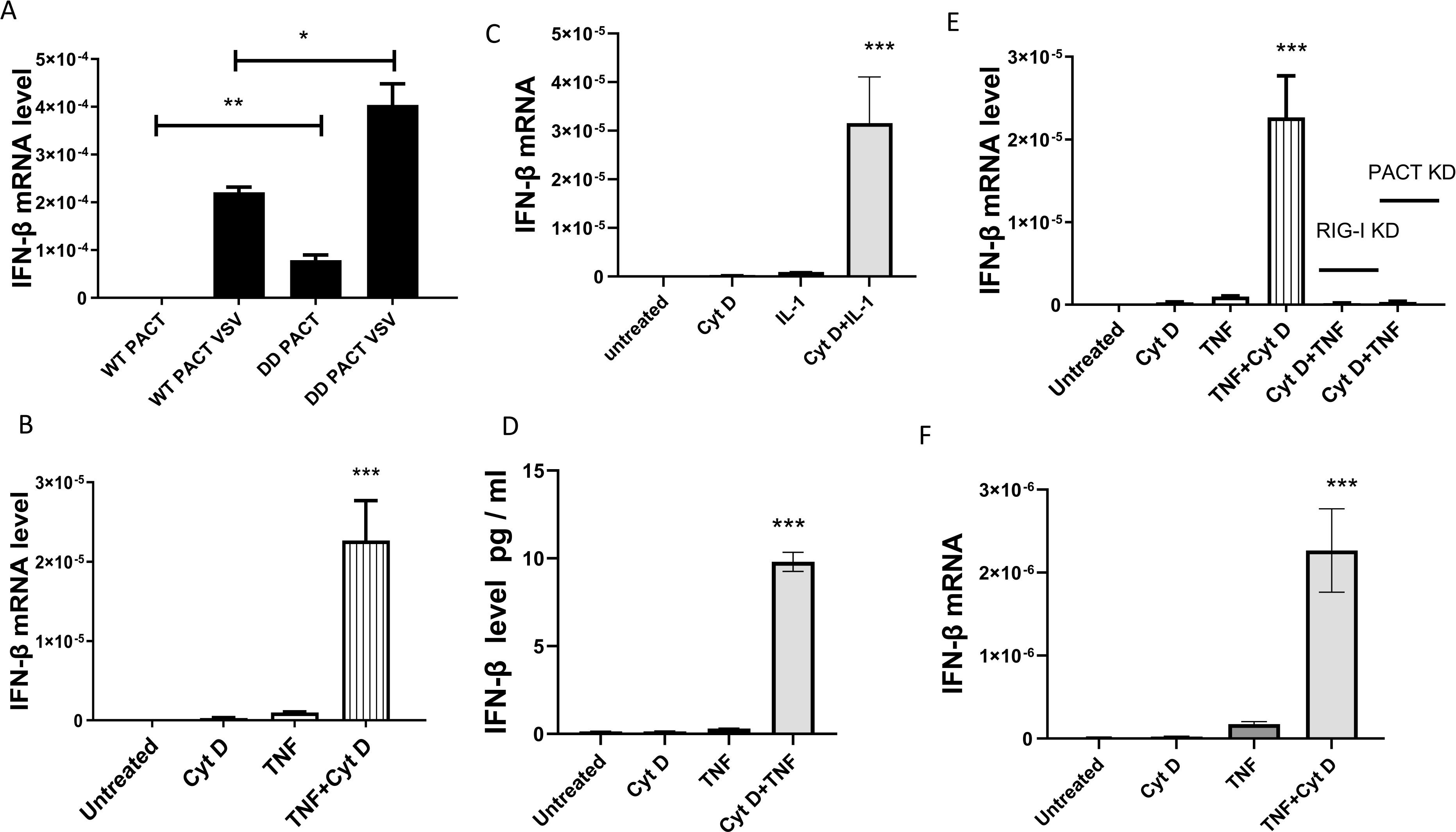
RIG-I and PACT mediated IFN-b induction by TNF-a or IL-1b. **(A)** WT or double phosphomimetic DD mutant of PACT was transfected to PACT-/- AML-12 cells. IFN-β mRNA expression was analyzed 8 hours post VSV infection. **(B)** AML-12 cells were treated with cytochalasin D for 1 hour followed by TNF-α treatment. Only cytochalasin D or TNF-α treated cells were used as controls. IFN-β expression was analyzed 8 hours post TNF-α treatment. **(C)** AML-12 cells were treated with cytochalasin D for 1 hour followed by IL-1β treatment. Only cytochalasin D or IL-1 treated cells were used as controls. IFN-β mRNA expression was analyzed 8 hours post IL-1 treatment. **(D)** AML-12 cells were treated with cytochalasin D for 1 hour followed by TNF-α treatment. Only cytochalasin D or TNF-α treated cells were used as control. IFN-β protein secreted to culture supernatant was quantified 12 hours post TNF-α treatment using ELISA. **(E)** WT AML-12 cells, PACT -/- and RIG-I -/- AML12 cells were treated with cytochalasin D for 1 hour followed by TNF-α treatment. Only cytochalasin D or TNF-α treated cells were used as controls. IFN-β mRNA expression was analyzed 8 hours post TNF-α treatment. **(F)** Mouse primary hepatocytes were treated with cytochalasin D for 1 hour followed by TNF-α treatment. Only cytochalasin D or TNF-α treated cells were used as controls. IFN-β mRNA expression was analyzed 8 hours post TNF-α treatment. (**A, B, C, D, E, F**: Mean ± SEM, N=3). *P<0.05, **P<0.01, ***P<0.001.

To explore the mechanism of PACT-mediated activation of RIG-I signaling, we examined physical interactions between the two proteins. Co-immunoprecipitation assays revealed a strong interaction between the two proteins in cells treated with Cyt D and TNF-α, but not TNF-α alone; a mild interaction was also observed in cells treated with Cyt D alone. These results indicated that RIG-I priming was essential for this interaction (Fig 2A). The above conclusion was confirmed by a cell-based assay, the proximal ligation assay (PLA). We recorded strong signals from cells treated with Cyt D and TNF-α, but not from untreated cells or cells treated with TNF-α or Cyt D alone (Fig 2B). Activation of RIG-I signaling by viral RNA is initiated by their interaction followed by recruitment of the enzyme TRIM25 to the signaling complex and K-63 ubiquitination of RIG-I; RIG-I activation by PACT followed the same pattern. TRIM25 was co-immunoprecipitated with RIG-I from cells treated with Cyt D and TNF-α (Fig 2C) and RIG-I was K-63 ubiquitinated in cells treated with Cyt D and TNF-α (Fig 2D) or Cyt D and IL-1β (Fig 2E). Similar K-63 ubiquitination was observed in Cyt D and TNF-α treated human liver HepG2 cells (Fig 2F) and mouse primary hepatocytes (Fig 2G).

**Figure 2.**
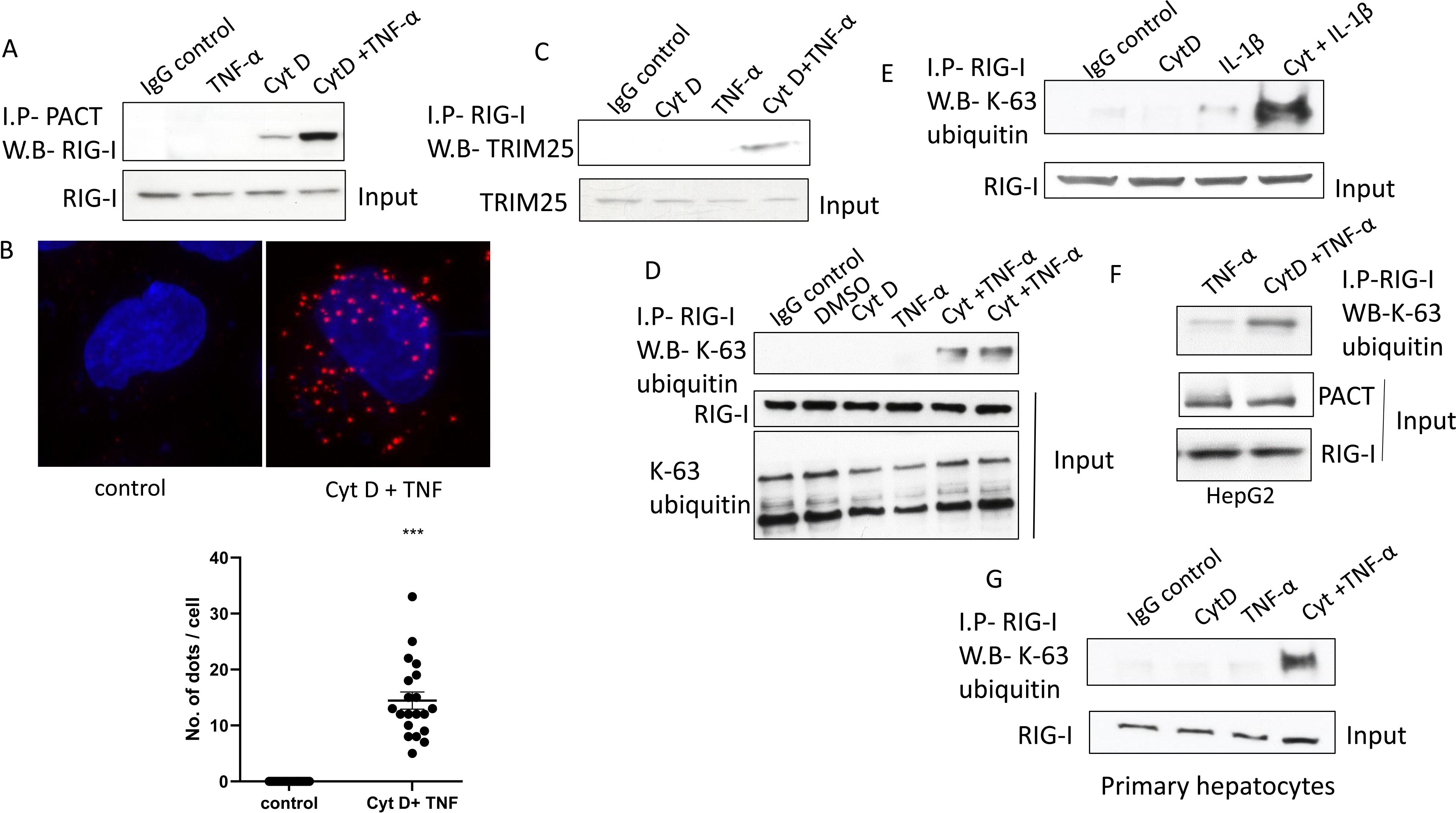
In TNF-a treated cells, PACT binds to RIG-I and triggers its ubiquitination. **(A)** AML-12 cells were treated with cytochalasin D for 1 hour followed by TNF-α treatment. Only cytochalasin D or TNF-α treated cells were used as control. The cells were harvested one hour post TNF-α treatment and PACT was immunoprecipitated from cell extracts. Co-precipitation of RIG-I was detected by Western blot analysis. **(B)** AML-12 cells were grown on cover slips and treated with cytochalasin D for 1 hour followed by TNF-α treatment. The cells were fixed by paraformaldehyde and stained with RIG- and PACT specific antibodies to perform PLA assays. Confocal images were acquired to demonstrate interaction between PACT and RIG-I. The graph below depicts the quantification of PLA dots in 20 cells. **(C)** AML-12 cells were treated with cytochalasin D for 1 hour followed by TNF-α treatment. Only cytochalasin D or TNF-α treated cells were used as controls. The cells were harvested one hour post TNF-α treatment, RIG-I was immunoprecipitated from the cell extracts and analyzed by Western blots for TRIM-25. **(D)** AML-12 cells were treated with cytochalasin D for 1 hour followed by TNF-α treatment. Only cytochalasin D or TNF-α treated cells were used as controls. The cells were harvested 1 hour or 2-hour post TNF-α treatment. RIG-I was immunoprecipitated from the cell extracts and analyzed by Western blot using K-63 Ub antibody. **(E)** AML-12 cells were treated with cytochalasin D for 1 hour followed by IL-1β treatment. Only cytochalasin D or IL-1β treated cells were used as controls. The cells were harvested 2-hour post IL-1β treatment, RIG-I was immunoprecipitated from the cell extracts and analyzed by Western blot using K-63 Ub antibody. **(F)** Human HepG2 cells were treated with cytochalasin D for 1 hour followed by TNF-α treatment. K-63 ubiquitination of RIG-I was analyzed as in **E**. **(G)** Mouse primary hepatocytes were treated with cytochalasin D for 1 hour followed by TNF-α treatment. Only cytochalasin D or TNF-α treated cells were used as controls. K-63 ubiquitination of RIG-I was analyzed as in **E**.

It is known that PACT is activated by phosphorylation of S287 in response to extracellular stress (20), but the relevant protein kinase has not been identified. We screened for the effects of inhibitors of various MAP kinases, activated by TNF-α or IL-1β, on IFN-β mRNA induction and identified p38 as the critical enzyme. A p38 inhibitor blocked both IFN-β mRNA induction (Fig 3A) and RIG-I K-63 ubiquitination (Fig 3B). As expected, PKR activation by PACT was also blocked by the p38 inhibitor (Fig 3C) and TNF-α or TPA treatment led to p38 phosphorylation (Fig 3D). To investigate whether PACT is the direct substrate of p38, we compared the amino acid sequences around the phosphorylation sites of PACT and those of ELK-1 and ATFII, two known substrates of p38 (21). Both the phospho acceptor sites and the putative p38 binding F sites in the three proteins were similar (Fig 3E), indicating that PACT might be a direct substrate of p38. To test this possibility, we mutated the F site residues of PACT and tested the effect on its functions. The mutant PACT did not support IFN-β mRNA induction (Fig 3F) or K-63 ubiquitination of RIG-I (Fig 3G), indicating that cytokine-activated p38 directly phosphorylates S287 of PACT.

**Figure 3.**
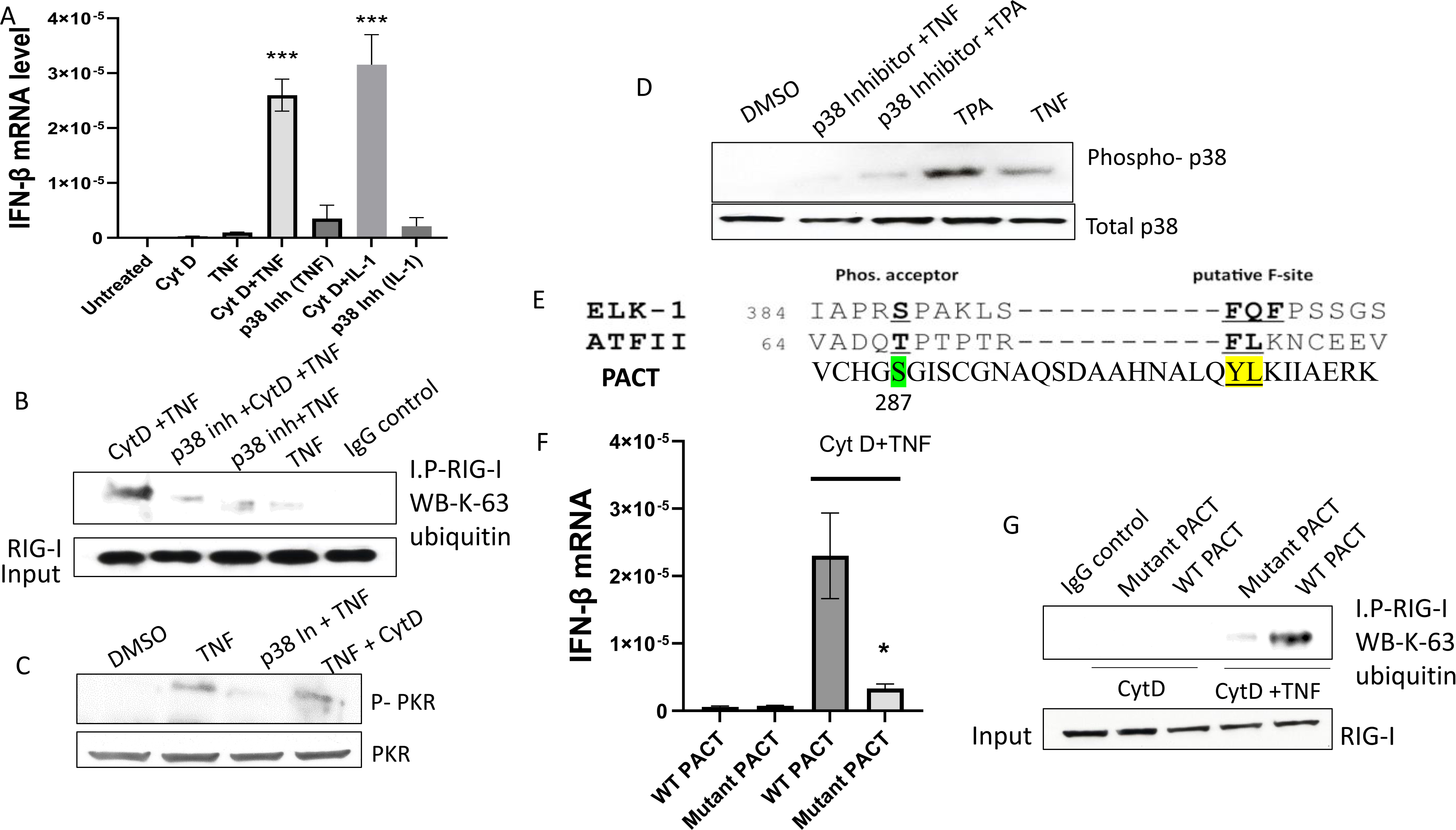
PACT is activated by direct phosphorylation by p38 in TNF-a or IL-1B treated cells. **(A)** Where indicated, AML-12 cells were pre-treated with the p38 inhibitor for 1 hour followed by cytochalasin D and cytokine treatment. IFN-β mRNA expression was analyzed 8 hours post treatment. **(B)** AML-12 cells were treated as in **A.** The cells were harvested one hour post TNF-α treatment, RIG-I was immunoprecipitated from the cell extracts and analyzed for K-63 ubiquitination. **(C)** AML-12 cells were pre-treated with p38 inhibitor or DMSO followed by TNF-α or cytochalasin D + TNF-α treatment. The cells were harvested 1 hour post TNF-α treatment and PKR phosphorylation was analyzed by western blotting. **(D)** AML-12 cells were pre-treated with p38 inhibitor or DMSO followed by TNF-α or TPA (12-O-Tetradecanoylphorbol-13-acetate) treatment. The cells were harvested 1 hour post TNF-α / TPA treatment and p38 MAPK phosphorylation was analyzed by western blotting. **(E)** The amino acid sequence of mouse PACT highlighting putative p38 MAPK binding site (F-site) compared with those of two known substrates of p38. These F-sites downstream to main phospho-acceptor serine residue are considered as the binding sites for p38 kinases. **(F)** WT or p38 mutant PACT expressing cells were treated with cytochalasin D for 1 hour followed by TNF-α treatment. IFN-β mRNA expression was analyzed 8 hours post TNF-α treatment. **(G)** WT and p38 mutant PACT expressing cells were treated with cytochalasin D for 1 hour followed by TNF-α treatment for 1 hour. RIG-I was immunoprecipitated from the cell extracts and analyzed for K-63 ubiquitination. (**A, F**: Mean ± SEM, N=3). *P<0.05, **P<0.01, ***P<0.001.

The observations, reported here, indicate that IFN-β can be induced by the RLR pathway without any participation of viral or cellular RNA. Such induction requires PACT to be activated by p38, a MAPK, that is activated by many extracellular stimuli including inflammatory cytokines. But RIG-I could bind to PACT only if it was primed. Both proteins needed specific modifications before their interaction; PACT needed to be phosphorylated and RIG-I needed to be dephosphorylated (22, 23). Further investigations should reveal the nature of physiological cues that lead to RIG-I priming. PACT activation by p38, on the other hand, can happen in response to a variety of extracellular stresses as observed for PKR activation by PACT. The structural features of PACT required for RIG-I activation were the same as those required for PKR activation, indicating that mutants of PACT which cannot bind dsRNA but still activate PKR (14), will be able to activate primed RIG-I as well. It is possible that in many cell-lines, PACT co-operates with viral RNA to induce IFN in infected cells. But, in some cell lines, such as AML-12, viral RNA could not induce IFN in the absence of PACT indicating that in such situations, virus infection primes RIG-I but PACT is the activating ligand (18)

## ACKNOWLEDGEMENTS

We would like to acknowledge Laura Nagy (Cleveland Clinic) for providing mouse primary hepatocytes for our experiments. This work was supported by the National Institutes of Health grants CA062220 and CA068782.

---

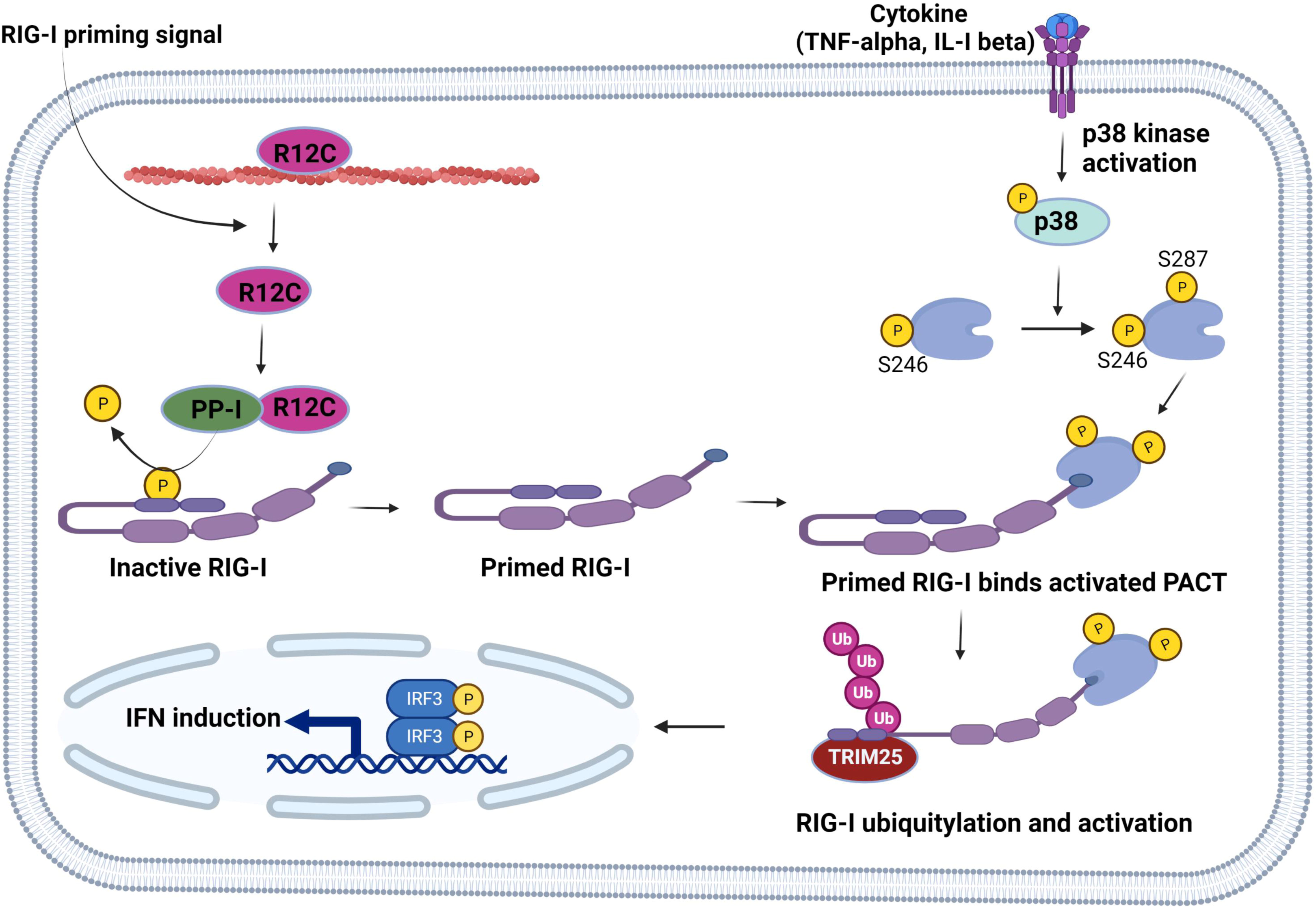

